# Kinetic and thermodynamic modeling of a voltage-gated sodium channel

**DOI:** 10.1101/2021.09.25.461780

**Authors:** Mara Almog, Nurit Degani-Katzav, Alon Korgnreen

## Abstract

Like all biological and chemical reactions, ion channel kinetics are highly sensitive to changes in temperature. Therefore, it is prudent to investigate channel dynamics at physiological temperatures. However, most ion channel investigations are performed at room temperature due to practical considerations, such as recording stability and technical limitations. This problem is especially severe for the fast voltage-gated sodium channel, whose activation kinetics are faster than the time constant of the standard patch-clamp amplifier at physiological temperatures. Thus, biologically detailed simulations of the action potential generation evenly scale the kinetic models of voltage-gated channels acquired at room temperature. To quantitatively study voltage-gated sodium channels’ temperature sensitivity, we recorded sodium currents from nucleated patches extracted from the rat’s layer five neocortical pyramidal neurons at several temperatures from 13.5 to 30oC. We use these recordings to model the kinetics of the voltage-gated sodium channel as a function of temperature. We show that the temperature dependence of activation differs from that of inactivation. Furthermore, we show that the sustained current has a different temperature dependence than the fast current. Our kinetic and thermodynamic analysis of the current provided a numerical model spanning the entire temperature range. This model reproduced vital features of channel activation and inactivation. Furthermore, the model also reproduced action potential dependence on temperature. Thus, we provide an essential building block for the generation of biologically detailed models of cortical neurons.

## Introduction

Generating biologically detailed models of neuronal excitability is a significant stepping stone on the path to simulating the function of the nervous system (Gold *et al.*, 2007; Keren *et al.*, 2009; Hay *et al.*, 2011; Reimann *et al.*, 2013; Almog & Korngreen, 2014, 2016; Markram *et al.*, 2015). Such models of neuronal excitability contributed important information within the scope of the study for which they were created. However, detailed models contain many crude assumptions and poorly constrained parameters (Almog & Korngreen, 2016). Thus, current biologically detailed models are ad hoc, unrealistic models functioning poorly once they are stretched beyond the specific problems for which they were designed (Almog & Korngreen, 2016). Thus, they are not appropriate for general reuse to address other scientific questions. Furthermore, due to their nature and their less-than-realistic virtual axons, which they include, it is hard to determine their validity within the context of the neuronal network.

There are many ways in which biologically detailed models can be improved. One of them is to extract and verify cellular parameters, thus reducing the overall uncertainty of the model. A prominent parameter set, extracted using standard techniques, is the kinetics of voltage-gated channels and their temperature dependence. Chemical and biological reactions are all modulated by temperature. In physiology, many processes’ rate increase with temperature, including electrical excitability (Hodgkin & Katz, 1949; Weight & Erulkar, 1976; Kim & Connors, 2012). Studies have shown that temperature-derived electrical excitability changes are primarily due to changes in transition rates controlling ion channel function and changes in ion channel conductance (Frankenhaeuser & Moore, 1963; Korogod & Demianenko, 2017). One of the highly influenced channels by changes to temperature is the voltage-gated sodium channel (Collins & Rojas, 1982; Rosen, 2001; Egri & Ruben, 2012), responsible for action potential generation and propagation. Thus, understanding the impact of temperature on voltage-gated sodium channels is vital for modeling neuronal excitability at physiological temperatures. Moreover, understanding how temperature affects action potentials is crucial for the detailed modeling of neuronal excitability (Collins & Rojas, 1982; Korogod & Demianenko, 2017; DeMaegd & Stein, 2020).

All biologically detailed models of neuronal excitability use kinetics of voltage-gated sodium channels recorded at room temperature. The lack of detailed kinetic analysis for the voltage-gated sodium channel at physiological temperature is primarily due to the instability of membrane patches at these temperatures and patch-clamp amplifiers’ limited ability to follow the channel’s rapid activation. Thus, the channel’s kinetics and conductance are adjusted using the temperature coefficient (Q_10_) in many neuronal excitability simulations. Classically Q_10_ was defined as the unitless ratio of the reaction rates separated by ten degrees. The reaction rates and conductances of the channels are exponentially scaled using this coefficient. A fixed value of Q_10_ is used (a value of 2.3 is prevalent in many simulations serving as a global temperature fudge factor). However, Q_10_ varies between channel types and is only quantitatively available for a small subset of channels (Decoursey & Cherny, 1998; DeMaegd & Stein, 2020). Furthermore, Q_10_ changes as a function of temperature (Collins & Rojas, 1982; Allen & Mikala, 1998; Thomas *et al.*, 2009) and membrane potential (Thomas *et al.*, 2009; Ranjan *et al.*, 2019).

Here we overcome the technical difficulties of recording voltage-gated sodium channel kinetics at physiological temperatures by recording at several lower temperatures and extrapolating the rate constants to 37°C. We show that the temperature dependence of activation differs from that of inactivation. Furthermore, we show that the sustained current has a different temperature dependence than the fast current. Our kinetic and thermodynamic analysis of the current provided a numerical model spanning the entire temperature range. This model reproduced vital features of channel kinetics and action potential generation.

## Methods

### Animals

All procedures were approved and supervised by Bar-Ilan university’s Animal Care and Use Committee and were in accordance with the National Institutes of Health Guide for the Care and Use of Laboratory Animals and the University’s Guidelines for the Use and Care of Laboratory Animals in Research. This study was approved by the National Committee for Experiments in Laboratory Animals at the Ministry of Health.

### Slice preparation

Slices (sagittal, 300 μm thick) were prepared from the somatosensory cortex of 12-16 days old Wistar rats that were killed by rapid decapitation using previously described techniques (Korngreen & Sakmann, 2000; Bar-Yehuda *et al.*, 2008; Almog & Korngreen, 2014; Almog *et al.*, 2018). Slices were maintained at room temperature in a chamber with an oxygenated artificial cerebrospinal fluid (ACSF) containing (mM): 125 NaCl, 25 NaHCO_3_, 2.5 KCl, 1.25 NaH_2_PO_4_, 1 MgCl_2_, 2 CaCl_2_, 25 Glucose, 0.5 Ascorbate (pH 7.4 with 5% CO_2_, 310 mosmol/kg). Pyramidal neurons from L5 in the somatosensory cortex were visually identified using infrared differential interference contrast (IR-DIC) videomicroscopy (Stuart *et al.*, 1993).

### Electrophysiology

Nucleated outside-out patches (Sather *et al.*, 1992; Korngreen & Sakmann, 2000; Almog & Korngreen, 2014) were extracted from the soma of L5 pyramidal neurons. Briefly, negative pressure (180-230 mbar) was applied when recording in the whole cell configuration, and the pipette was slowly retracted. Provided that the retraction was gentle it was possible to obtain large patches of membrane engulfing the nucleus of the neuron. After the extraction of the patch, the pressure was reduced to 30-40 mbar for the duration of the experiment. All measurements from nucleated patches were carried out with the Axopatch-700B amplifier (Axon Instruments, Foster City, CA) using a sampling frequency of 100 kHz and filtered at 20 kHz. The capacitive compensation circuit of the amplifier was used to reduce capacitate transients. Nucleated patches were held at −60 mV. Leak was subtracted using a P/4 online protocol applied at a negative holding potential of −100 mV. We only analyzed currents from which the linear leak was cleanly subtracted by the P/4 protocol, thus leaving no residual of the capacitance transient. This was quite straightforward when the capacitance transient did not saturate the amplifier or exceed the dynamic range of the A/D converter. Due to the small membrane surface the total capacitance of a nucleated patch was small and allowed for excellent temporal control of the voltage-clamp amplifier. We verified this by measuring the decay of the uncompensated capacitance transients that decayed to baseline in under 40 μs. To be on the safe side, we removed the first 50 μs from the data before analysis. Note that this may result in an under- estimation of the channel kinetics. Patch pipettes (4-7 MΩ) were coated with Sylgard (DOW Corning). To record sodium currents in nucleated patches the pipettes were filled with a solution containing 120 mM Cs-gluconate, 20 mM CsCl, 10 mM HEPES, 4 mM MgATP, 10 mM phosphocreatine, 1 mM EGTA, 0.3 mM GTP (pH=7.2, CsOH). The liquid junction potential of −11 mV generated by this pipette solution was not corrected for during the experiments or offline analysis. In current clamp experiments (Fig. 11) the pipette solution contained (mM): 125 K-gluconate, 20 KCl, 10 HEPES, 4 MgATP, 10 Na-phosphocreatin, 0.5 EGTA, 0.3 GTP and 0.2 % biocytin (pH=7.2 with KOH). Bath temperature was controlled by a custom built Peltier device that controlled the temperature of the ACSF flowing into the recording chamber. The temperature next to the slice was monitored by a thin thermocouple and the temperature of the Peltier device was changed to ensure the target temperature. Experiments were carried out at four temperature ranges as detailed in the following table. In the figures and text we refer to the median temperature of these sets.

### Data analysis and numerical simulations

All off-line data analyses including curve fitting were carried out with IgorPro (WaveMetrics, Lake Oswego, USA). Experimental results were observed in cells from two or more animals. It is customary in many papers to display representative or typical raw traces. This practice might impose a human selection bias inadvertently selecting unusually nice recordings. To avoid this, we present the population average of all the experiments that were analyzed. The standard deviation of each data point could thus be calculated. This also has the advantage of averaging inter-patch variability in channel density.

The analysis presented in this paper contained several data reduction stages. First, channel activation and inactivation parameters were estimated using a curve fit to current traces. Then rate constants were calculated from these parameters. Finally, curve fitting extracted the functional dependence of the rate constants on voltage and temperature. This multistage analysis required consideration of error propagation between analysis stages. In general, we calculated the propagated error using the partial derivative method. Thus, for a general function f(x,y), the propagated error Δ*f* was approximated as:

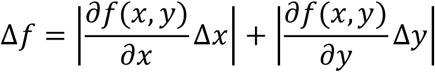

Where Δ*x* and Δ*y* are the experimentally measured standard deviations of x and y respectively. Curve fitting was performed using IgorPro’s intrinsic fitting algorithm and multiple curve fitting was performed with the GlobalFit package of IgorPro. To estimate the error of each fitted parameter we repeated this fitting procedure on data sets sampled with replacements (Bootstrap).

Simulations of ionic currents and action potentials were programmed using NEURON 7.3 (Carnevale & Hines, 2006). All simulations were performed with an integration interval of 5 μs to ensure a stable numerical solution of the differential equations. Ion channel models were implemented using the NMODL extension of NEURON (Hines & Carnevale, 2000).

## Results

We recorded sodium currents, presumably of Nav1.2 channels (Hu *et al.*, 2009), from nucleated patches using a Cesium-based pipette solution to block voltage-gated potassium currents. Fast inward currents were detected following a depolarizing voltage-clamp step from a holding potential of –110 mV (Fig. 1). We only analyzed data from patches displaying stable recordings, precise leak subtraction, and no current rundown. We applied a custom-built Peltier device to control the ACSF intake’s temperature to the bath surrounding the brain slice. The temperature of ACSF intake was adjusted until bath temperature, measured by a thermocouple, reached the desired value. As detailed in Table 1, we recorded at four median temperatures (13.5, 19.5, 25, and 30°C). To thoroughly compare the currents at different temperatures, we performed a point-by-point averaging of all recordings from a specific temperature. For example, the current recordings at 13.5°C displayed in Figure 1 are an average of 10 recordings from different patches. This global averaging procedure is not different than the standard analysis method in which the peak current at each voltage is averaged between patches to generate I-V curves. However, this procedure provided better visual representation of the data and a standard deviation for each point in the recordings. As expected, the peak sodium current increased as the median temperature increased from 13.5 to 30°C. Interestingly, the sustained current decreased as the temperature increased, which was manifested by the marked reduction in the tail current (Fig. 1. Open circles in the left panels). For each temperature, we measured the peak sodium current and plotted it as a function of the pipette potential (Fig. 1, right panels). These plots were fitted with:

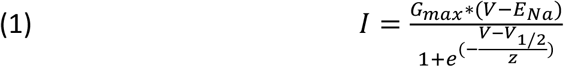

where *G*_*max*_ is the maximal conductance, *E*_*na*_ is the sodium reversal potential, *V*_*1/2*_ is the voltage of half activation, and *z* is the slope of the Boltzmann curve. The maximal conductance of the transient current increased approximately two fold between 13.5 and 30°C (Fig. 2A). Interestingly, the maximal conductance of the sustained current, calculated from the peak of the tail currents, decreased approximately two fold over this temperature range (Fig. 2A). Linear extrapolation of the change in the conductance of the sustained current to 37°C gave an estimate of 0.3 ±0.4 nS suggesting that at physiological temperatures the sustained current is small. In addition to the change in maximal conductance of the transient current, its voltage of half activation shifted to lower potentials (Fig. 2B) and the slope of the Boltzmann curve steepened (Fig. 2C) as a function of temperature.

**Figure 1:**
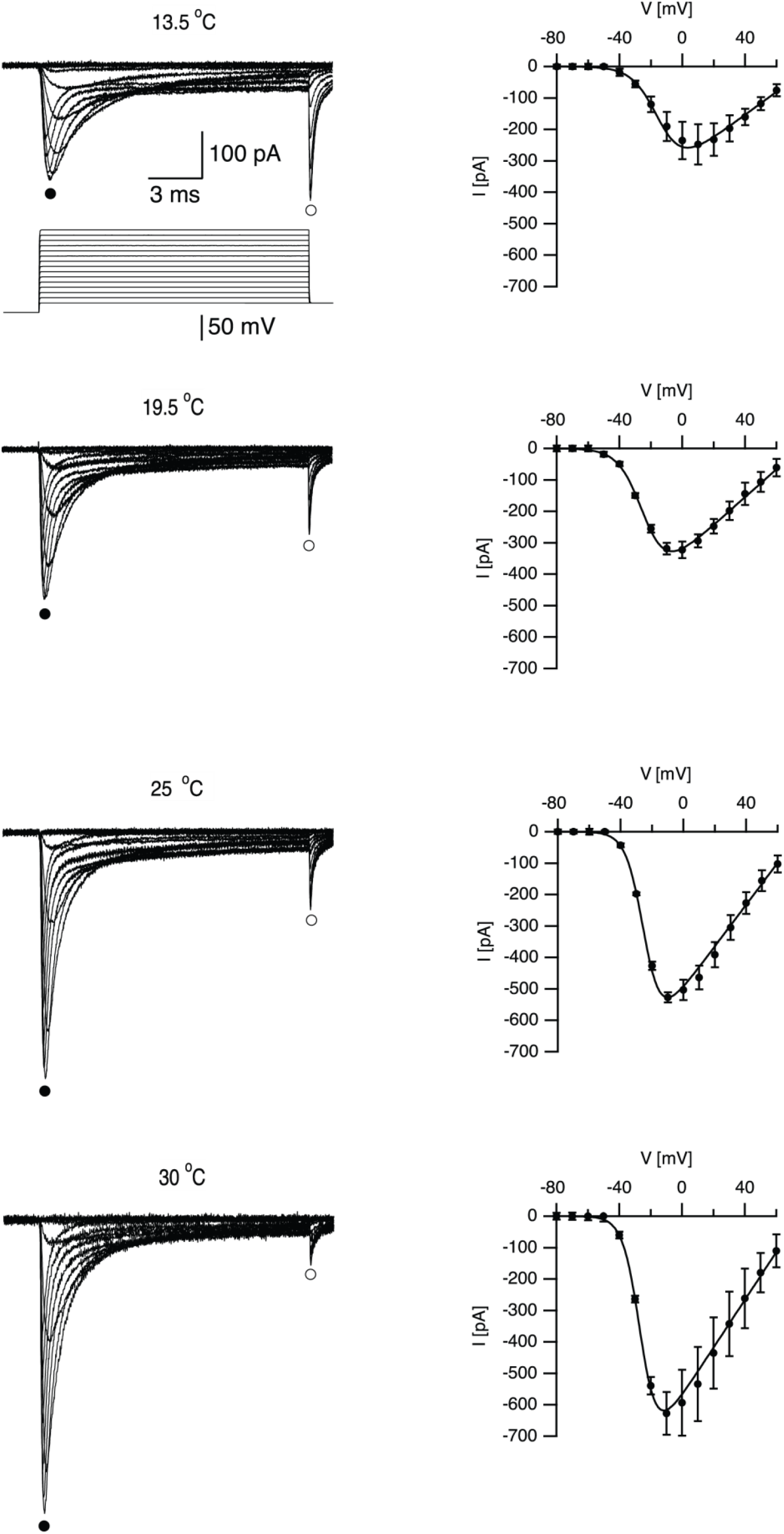
Temperature affects voltage-gated sodium channel activation. Population averages of sodium currents recorded at four temperatures are shown on the left column. The traces are averages of the raw recordings made from 10 (13.5°C), 8 (19.5°C), 7 (25°C), and 8 (30°C) recording sessions from different nucleated patches. The voltage-clamp protocol is displayed below the top traces. Following a 100 ms pre-pulse to −110 mV the membrane potential was depolarized to the test potential at 10 mV increments. Current-voltage curves, displayed on the right column, were calculated from the traces displayed on the left from the peak negative current recorded at each voltage (indicated by the filled circles on the left). Continuous lines in the current-voltage traces are the fit of the data to Eq. 1. Error bars are SEM.

**Figure 2:**
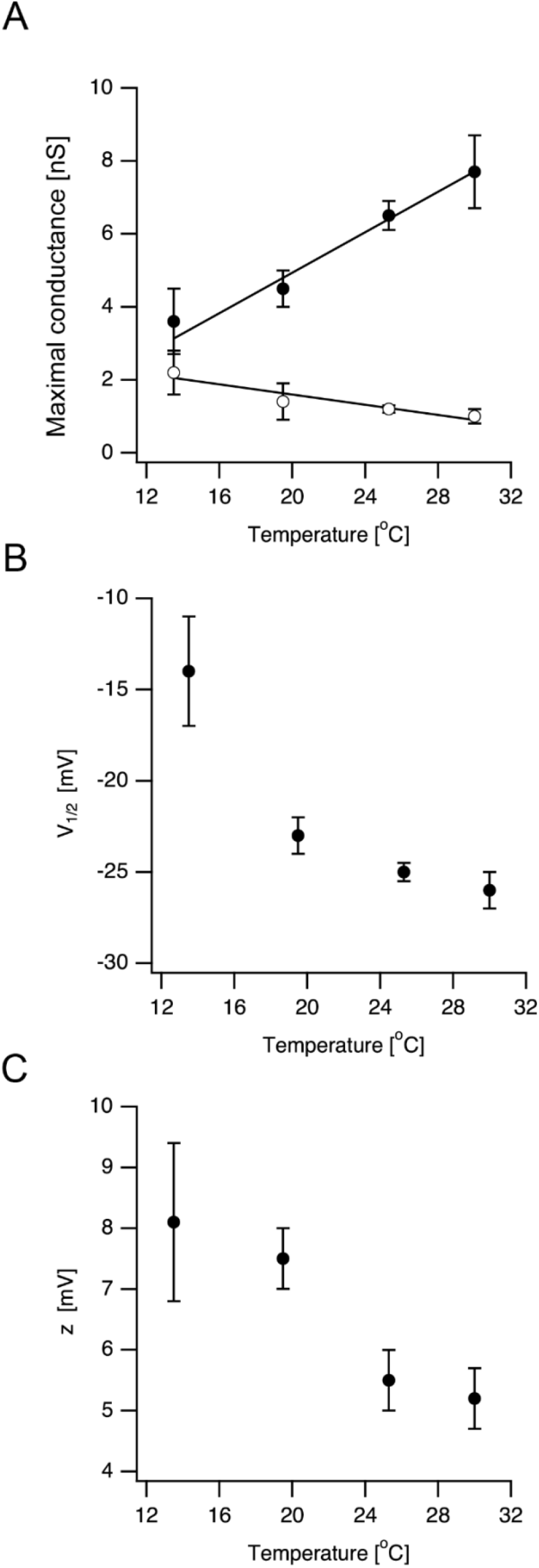
Dependence of voltage-gated sodium channel activation on temperature. A. Maximal ion channel conductance (Gmax) calculated from the fit of the peak negative current displayed in Fig. 1 to Eq. 1 (filled circles) and from the peak tail current (open circles here and also noted in Fig. 1). Lines are linear regression fit of the data to a straight line. B. Voltage of half-activation (V_1/2_) plotted as a function of temperature. C. the slope of the activation curve (z) plotted as a function of temperature. All error bars are SEM.

**Table 1:** temperature ranges applied in this study. N is the number of patches recorded from at a specific temperature.

| Average Temperature (°C) | N | Standard deviation | Median Temperature (°C) |
| --- | --- | --- | --- |
| 13.7 | 44 | 0.4 | 13.5 |
| 19.4 | 42 | 0.7 | 19.5 |
| 25.4 | 96 | 1.0 | 25 |
| 30.1 | 24 | 1.6 | 30 |

Our goal was to provide a model for sodium-channel kinetics that will be functional at physiological temperatures. Since the sustained current decreased at higher temperatures, we decided to analyze only the fast component of the current. We have previously shown that the activation of the sodium current followed Cole-Moore kinetics (Almog *et al.*, 2018). We suggested that the full Markov chain model for the voltage-gated sodium channel should probably contain at least 7-8 closed states that lead to onset latency. We initially used genetic algorithms (Keren *et al.*, 2005; Gurkiewicz & Korngreen, 2007; Gurkiewicz *et al.*, 2011) to globally fit such Markov models to the data presented in Figure 1. However, accounting for both voltage and temperature dependence in these models caused parameter inflation (Almog & Korngreen, 2016) that hampered the ability of the fitting algorithm to reach a global minimum. Moreover, it is probable that, when recording macro currents from a multi-channel patch, it is possible to observe only the temperature dependence of the final closed-open transition and not that of the hidden closed-closed transitions. Therefore, we decided to model the channel kinetics using standard Hodgkin-Huxley, assuming a concerted closed-open transition-like model in order to capture the temperature dependence of the channel. Current traces were analyzed assuming the general model:

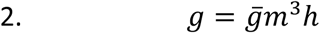

Where *g* is the conductance, 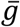 is the maximal conductance, *m* is the activation gate and *h* is the inactivation gate. Given the small contribution of the sustained current at physiological temperatures we analyzed only the transient current. Inward currents (Fig. 3A) generated by the activation protocol (Fig. 1) were fitted with:

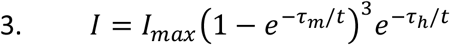

**Figure 3:**
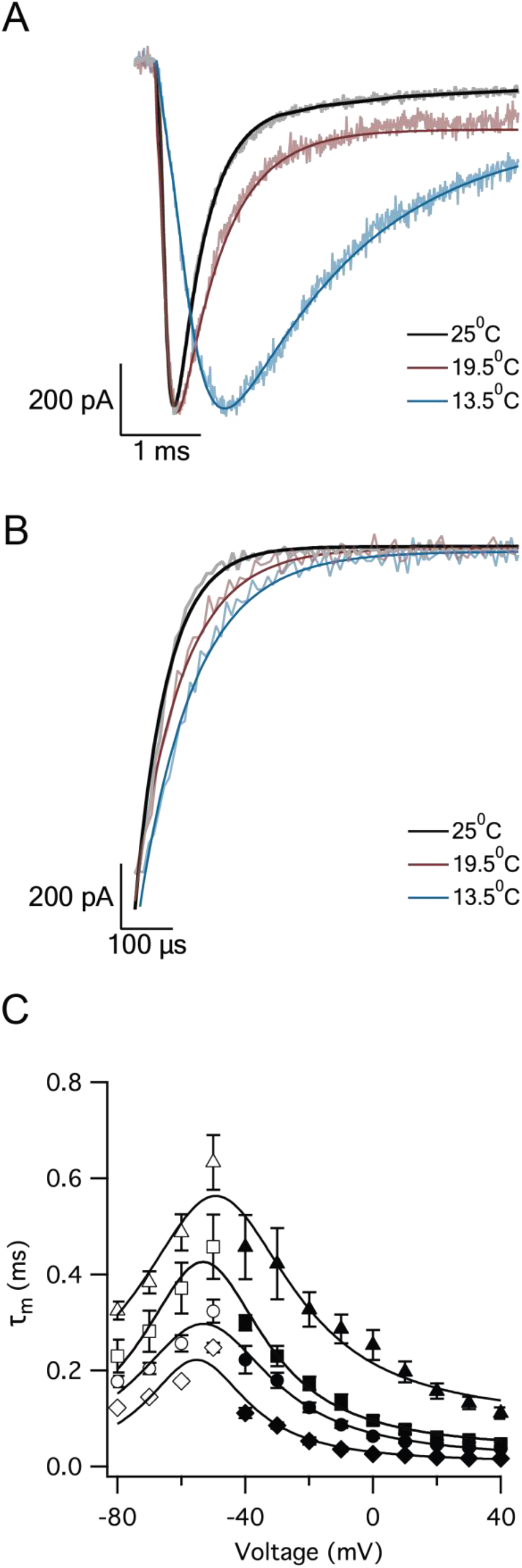
Kinetic analysis of channel activation. A. Representative curve fitting of Eq. 3 to sodium currents at several temperatures. B. representative curve fitting of Eq. 4 to sodium tail currents at several temperatures. C. activation time constants extracted from curve fitting of activation fitting (filled symbols) and deactivation fitting (empty symbols). The figure displays the averaged time constants for 13.5°C (triangles), 19.5°C (squares), 25°C (circles), and 30°C (diamonds). Continues lines are curve fitting of the data in each temperature to a Gaussian. Error bars are SEM.

Where I_max_ is the maximal current in each fitted sweep, τ_m_ is the activation time constant and τ_h_ is the inactivation time constant. Correspondingly, τ_m_ was estimated from current deactivation (tail current displayed in Fig. 3B) by fitting with:

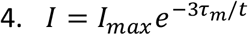

The time constant for activation and deactivation (Fig. 3C) displayed a characteristic asymmetric bell-shaped curve. As expected the time constant for activation decreased with temperature indicating faster reaction kinetics. To quantitatively measure the dependence of the activation and deactivation processes on temperature, we assumed that the rate constant dependence on temperature could be described by the Arrhenius equation (Arrhenius, 1889). For example the temperature dependence of the forward rate constant of the activation process was:

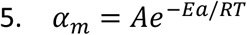

Where *α*_*m*_ is the forward rate constant, A is a proportionality constant, Ea is the activation energy, R is the gas constant and T is the absolute temperature. A similar expression, with the different activation energy, can be written for the backward rate constant *β*_*m*_. Since *τ*_*m*_ ≈ 1/*α*_*m*_ at depolarized potentials it was possible to use the Arrhenius plot (Fig. 4A) to roughly estimate *Ea* by plotting ln(*τ*) as a function of 1/T. We performed this analysis for membrane potentials above −20 mV (Fig. 4A) generating a family of roughly parallel linear plots. Averaging the slope of these curves estimated Ea to be 85 6 kJ/(mol·°K). Performing the same analysis for time constants derived from deactivating current traces (where *τ*_*m*_ ≈ 1/*β*_*m*_) provided an estimate for Ea of 42 1 kJ/(mol·°K) (Fig. 4B).

**Figure 4:**
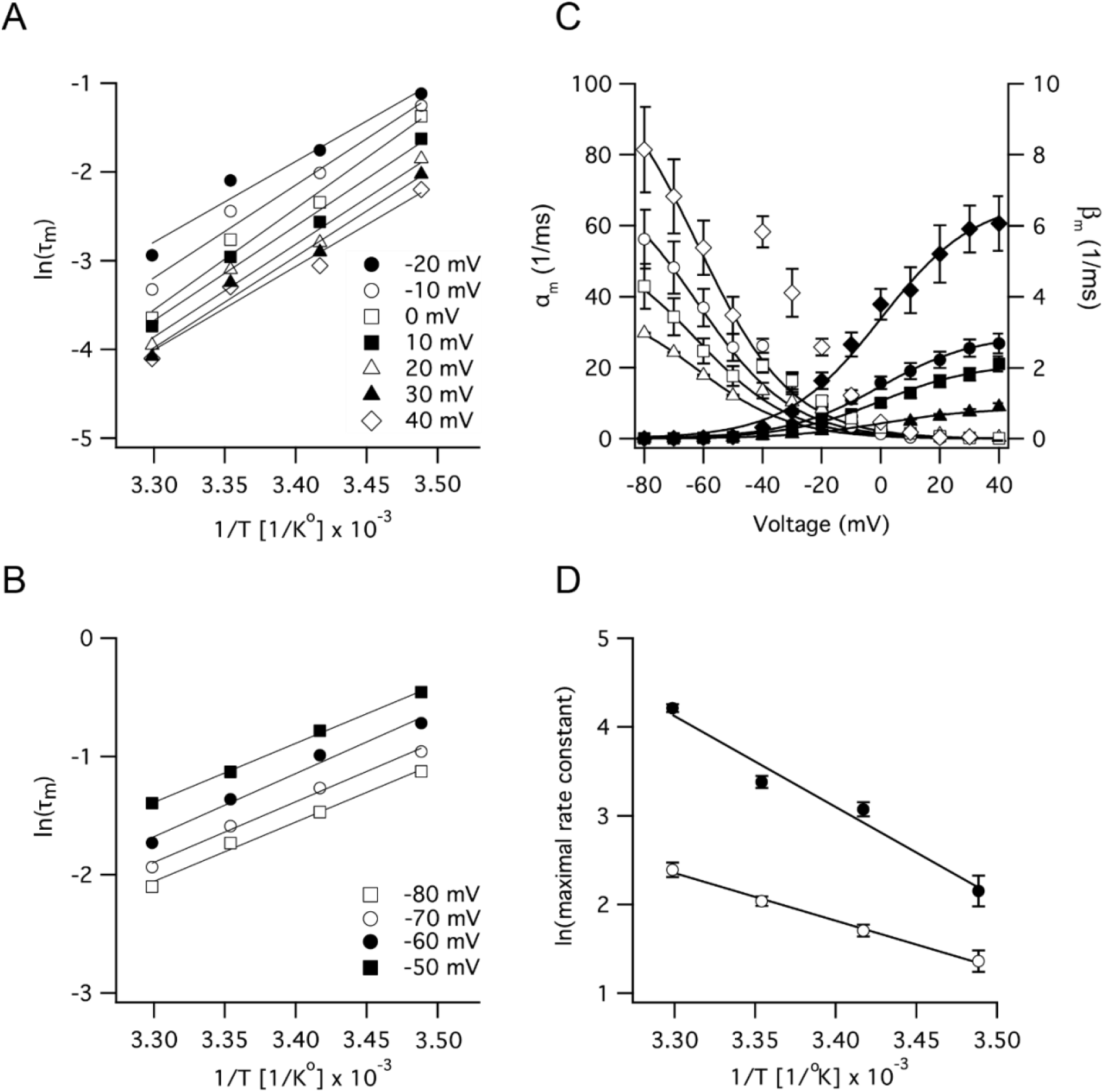
Thermodynamic analysis of channel activation. A. Arrhenius plots calculated from time constants derived from analysis of channel activation (Fig. 3A). Lines are linear regression fits to data from a specific voltage, as denoted in the legend. B. Arrhenius plots calculated from time constants derived from analysis of channel deactivation (Fig. 3B). C. Forward (filled symbols) and backward (empty symbols) rate constants derived using the Hodgkin-Huxley analysis from the data in Figures 1 and 3. Smooth lines are global curve fits of eq. 7 to the forward rate constant and eq. 8 to the backward rate constant. D. Arrhenius plots of the maximal rate constants obtained from the curve fit performed in C. Smooth lines are linear regression lines enabling calculation of the activation energies of the forward and backward transitions.

This thermodynamic analysis indicated that the forward and backward rate constants in the Hodgkin-Huxley model of the sodium channel are scaled with temperature according to standard rate theory (Stover *et al.*, 1974) and obey Eq. 5. This scaling suggested that these rate constants should display similar voltage dependence. Thus, we calculated *α*_*m*_ and *β*_*m*_ from the steady-state activation curves (Fig. 1) and time constants (Fig. 3C) and used global curve fitting (the Global Fit package of IgorPro) to fit them to a single, temperature scaled, voltage-dependent curve (Fig. 4C). The forward rate constant, *α*_*m*_, displayed a sigmoidal dependence on voltage (Fig. 4C) and therefore was fitted with the general equation:

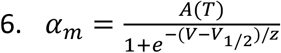

Where A is a temperature dependent scaling factor, V_1/2_ the voltage of half-activation, and z is the slope of the sigmoidal curve. In this global fit the values of z and V_1/2_ were forced to be the same for all *α*_*m*_ curves in Figure 4C while A(T) was estimated separately for each curve. This sigmoidal curve fitting estimated z to be 14.8 ± 0.6 mV and to be −3 ± 1 mV from all the data presented in this figure (parameter error estimation by bootstrap). The logarithm of A(T) was plotted as a function of 1/T which provided another estimate for Ea=85 ± 2 kJ/(mol·°K) (Fig. 4D) agreeing with the Arrhenius analysis from Figure 4A. Thus, the explicit equation for *α*_*m*_ was:

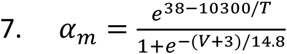

A similar fitting procedure was applied to *β*_*m*_ culminating with the equation:

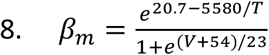

With z = −23 ± 4 mV and V_1/2_ = −54 ± 9 mV. The logarithm of A(T) was plotted as a function of 1/T which provided another estimate for Ea=44 ± 2 kJ/(mol·°K) (Fig. 4D) agreeing with the Arrhenius analysis from Figure 4B.

In agreement with the channel activation (Fig. 1), currents generated by a protocol designed to measure channel inactivation increased as the median temperature was increased from 13.5 to 30°C (Fig. 5). Inactivation curves (Fig. 5) were fitted with:

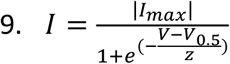

**Figure 5:**
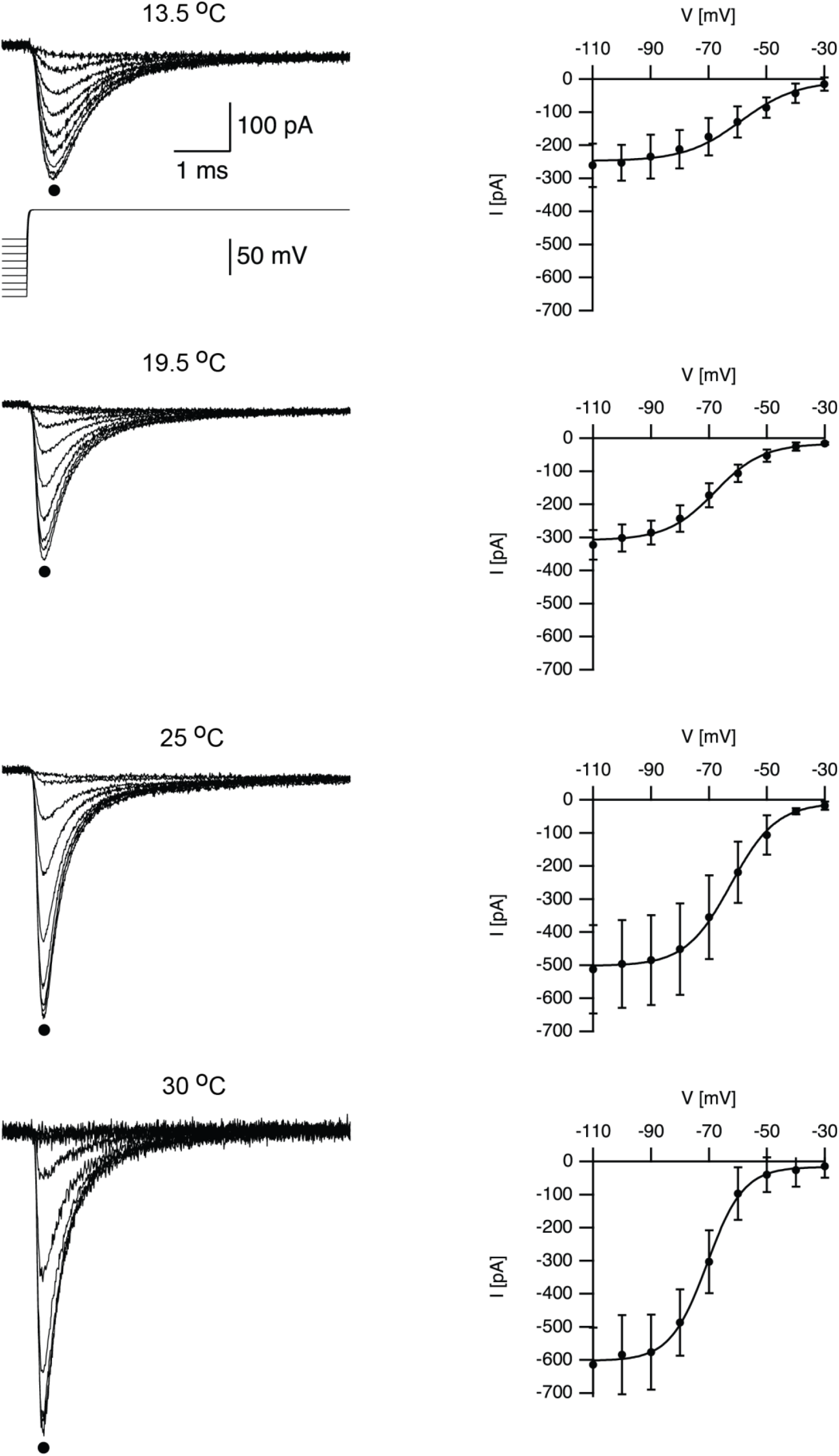
Temperature affects voltage-gated sodium channel inactivation. Population averages of sodium currents recorded at four temperatures are shown on the left column. The traces are averages of the raw recordings made from 7 (13.5°C), 8 (19.5°C), 15 (25°C), and 4 (30°C) recording sessions from different nucleated patches. The voltage-clamp protocol is displayed below the top traces. Following a 100 ms pre-pulse to several potentials at 10 mV increments the membrane potential was depolarized to the test potential of zero mV. Current-voltage curves, displayed on the right column, were calculated from the traces displayed on the left from the peak negative current recorded at each voltage (indicated by the filled circles on the left). Continuous lines in the current-voltage traces are the fit of the data to Eq. 9. Error bars are SEM.

Where *I*_*max*_ is the maximal patch current. The peak patch conductance extracted from this analysis recordings was linearly dependent on temperature (Fig. 6A) while the voltage of half activation (Fig. 6B) and the slope of the Boltzmann curve (Fig. 6C) did not display a clear dependence on temperature. The time constant of inactivation was estimated from Eq. 3 at potentials above activation threshold and from recovery from inactivation protocols below this threshold (Fig. 7). At 30°C it was not possible to hold patches for the long durations required for recovery protocols. Thus, the time constant for inactivation at this temperature was estimated only from activation protocols (Fig. 7D). Arrhenius analysis of the time constant over all the investigated voltage range (Fig. 8A) suggested, similarly to the analysis of the activation time constant (Fig. 4), that the inactivation reaction was exponentially dependent on temperature. However, we did not detect different activation energies over different voltage-ranges. The average activation energy derived from this analysis was 67 ± 8 kJ/(mol·°K).

**Figure 6:**
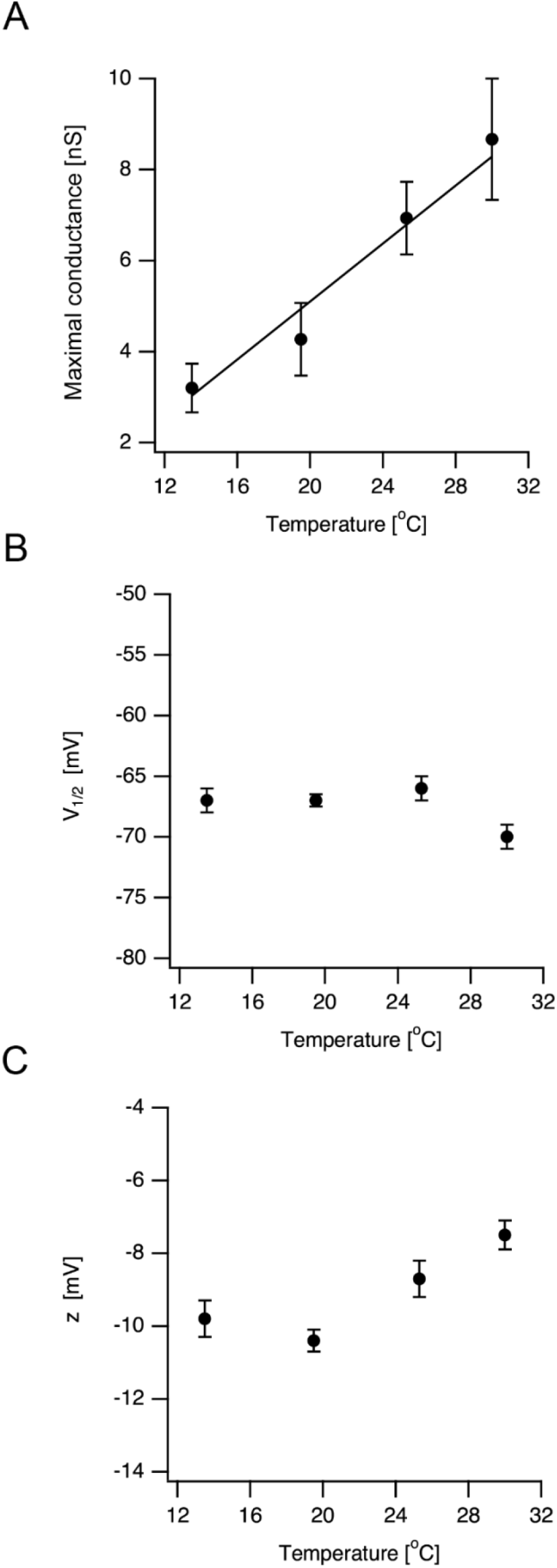
Dependence of voltage-gated sodium channel inactivation on temperature. A. Maximal ion channel conductance (Gmax) calculated from the fit of the peak negative current displayed in Fig. 6 to Eq. 1 (filled circles). The line is a linear regression fit of the data to a straight line. B. Voltage of half-inactivation (V_1/2_) plotted as a function of temperature. C. the slope of the inactivation curve (z) plotted as a function of temperature. All error bars are SEM.

**Figure 7:**
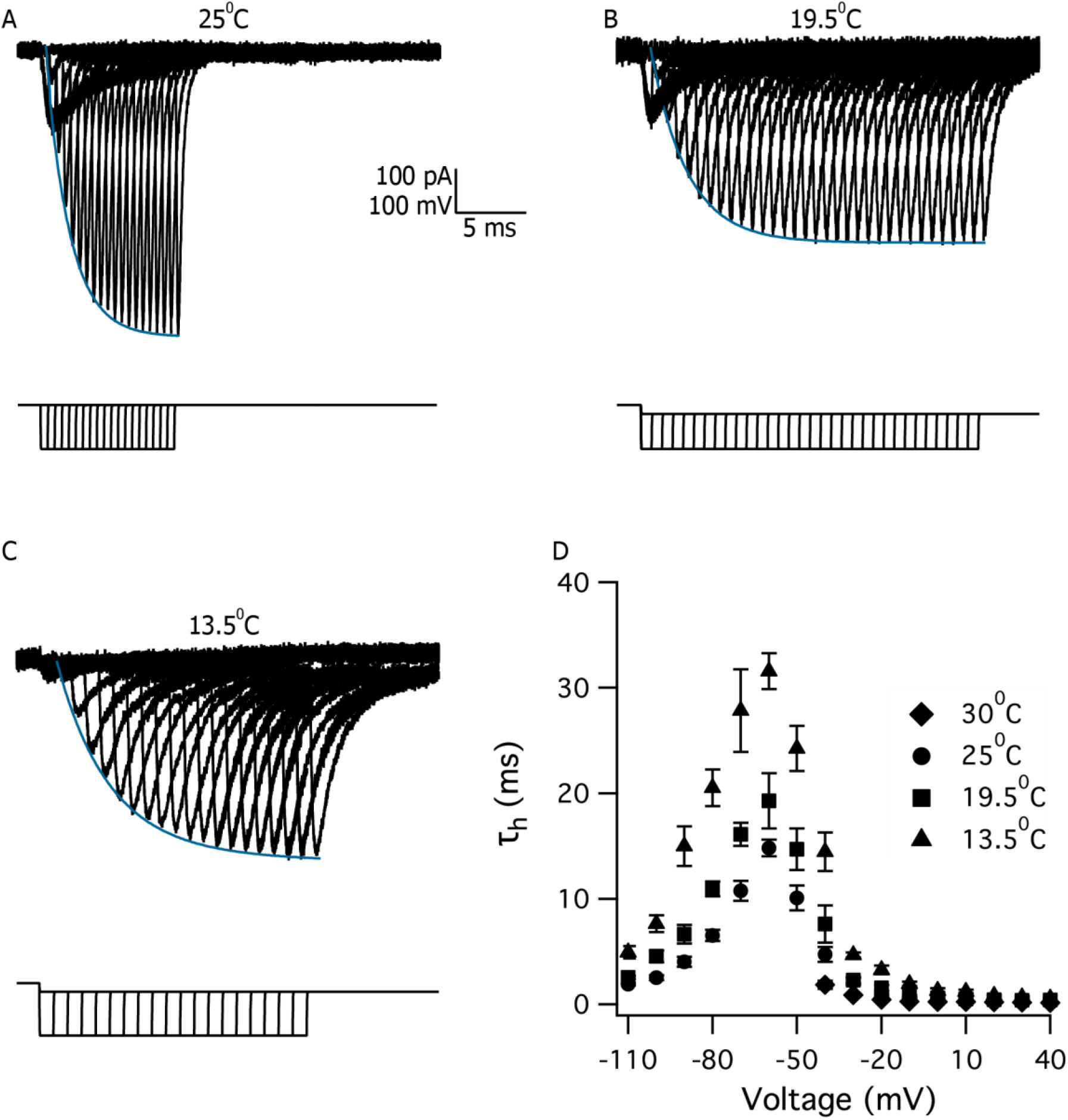
Kinetic analysis of channel inactivation. A. Recovery from inactivation recorded at 25°C using a triple pulse protocol. The membrane potential was clamped to zero mV for 50 ms to completely inactivate the channels (truncated from the figure), then the membrane was stepped to −100 mV for varying durations as shown below the current traces. Following this conditioning pulse the membrane potential was again stepped to zero mV to read the fraction of the current that recovered from inactivation. The smooth line is an exponential curve fit to the peaks of inward current. B. Similar to A save that the recording was performed at 19.5°C and the recovered current fraction was read using a voltage step to −10 mV. C. Similar to A save that the recording was performed at 13.5°C and the recovered current fraction was read using a voltage step to −10 mV. D. inactivation time constants extracted from curve fitting (Fig. 3A) and recovery from protocols displayed in A-C. The figure displays the averaged inactivation time constants for 13.5°C (triangles), 19.5°C (squares), 25°C (circles), and 30°C (diamonds). Error bars are SEM.

This scaling suggested that these rate constants should display similar voltage dependence. Thus, we calculated *α*_*h*_ and *β*_*h*_ from the steady-state inactivation curves (Fig. 5) and time constant (Fig. 7D) and used curve fitting to fit them to a single, temperature scaled, voltage-dependence (Fig. 8B). The forward rate constant, *α*_*h*_, displayed a dependence on voltage (Fig. 8C) and therefore was fitted with the general equation:

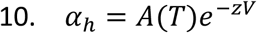

**Figure 8.**
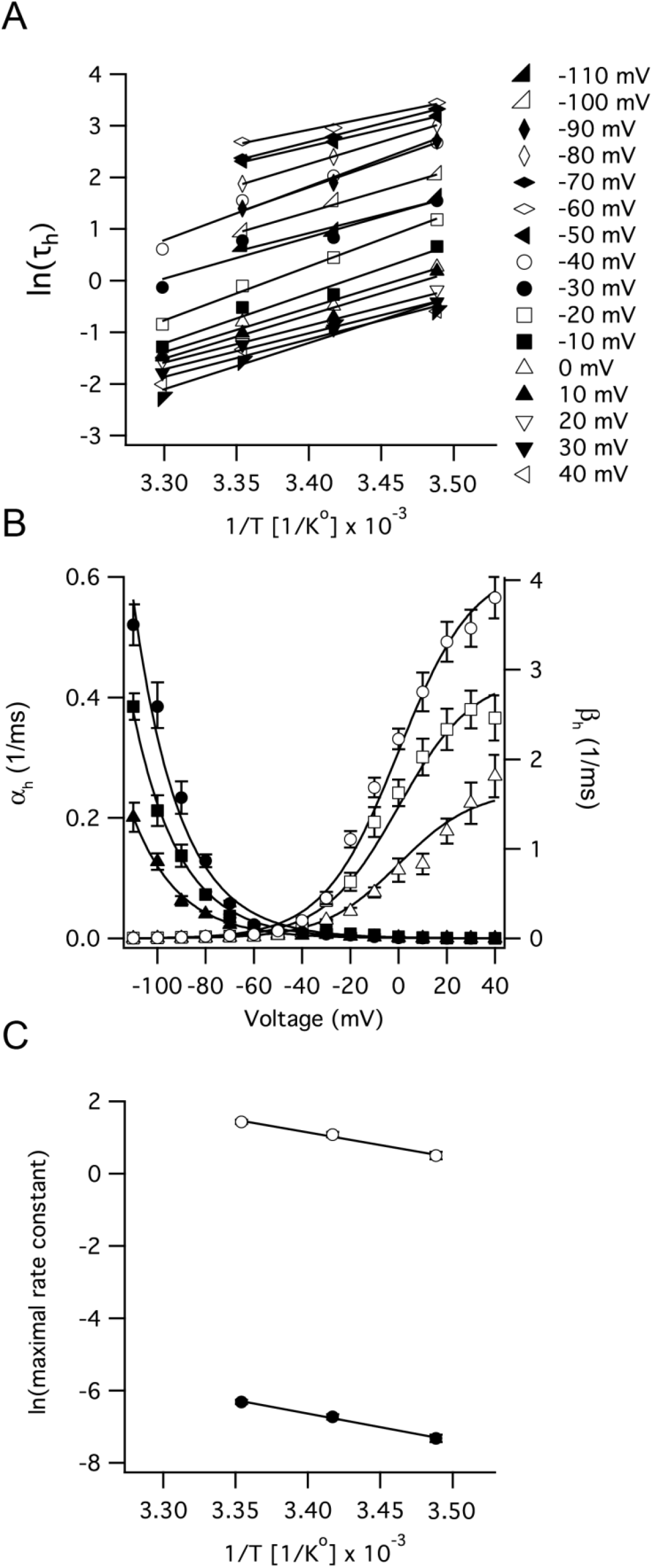
Thermodynamic analysis of sodium channel inactivation. A. Arrhenius plots calculated from time constants derived from analysis of channel inactivation (Fig. 7D). Lines are linear regression fits to data from a specific voltage, as denoted in the legend. B. Forward (filled symbols) and backward (empty symbols) rate constants derived using the Hodgkin-Huxley analysis from the data in Figures 5 and 7. Smooth lines are global curve fits of eq. 11 to the forward rate constant and eq. 12 to the backwardrate constant. C. Arrhenius plots of the maximal rate constants obtained from the curve fit performed in B. Smooth lines are linear regression lines enabling calculation of the activation energies of the forward and backward transitions.

Where A(T) is a temperature dependent scaling factor (signifying the maximal rate constant at a given temperature), and z is the rate of the exponential curve. In this two parameter fit the value of z was forced to be the same for all *α*_*h*_ curves in Figure 8B while A(T) was estimated separately for each curve. This exponential curve fitting estimated z to be 0.049 ± 0.001 mV^−1^. The logarithm of A(T) was plotted as a function of 1/T (Fig. 8C) which provided another estimate for Ea=62 ± 4 kJ/(mol·°K) agreeing with the Arrhenius analysis from Figure 8A. Thus, the equation for *α*_*m*_ was:

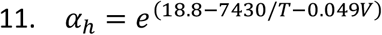

Using a sigmoidal curve fitting, similar to that applied to the activation rate constants, *β*_*h*_ was estimated to be:

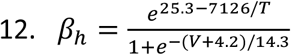

With z = 14.3 ± 0.8 mV and V_1/2_ = 4.2 ± 1.1 mV. The logarithm of A(T) was plotted as a function of 1/T which provided another estimate for Ea=58 ± 6 kJ/(mol·°K) (Fig. 8C) agreeing with the Arrhenius analysis from Figure 8A.

We simulated (Fig. 9), using NEURON (Carnevale & Hines, 2006) and the above quantitative description of the activation and inactivation of the sodium channel, voltage-clamp experiments replicating the traces displayed in Figures 1 and 5. The simulated currents deviated from the experimental ones in two ways. First, the prominent tail currents shown in Figure 1, were absent from the simulation. This is due to our decision not to simulate the sustained sodium current in this study. Second, the change in the peak amplitude of the sodium current was less prominent in the simulation (Fig. 9) than in the experiment (Figs. 1, 2A, 5, and 6A). This is probably due to the temperature dependence of the channel conductance (Milburn *et al.*, 1995) that was not simulated here. Next, we simulated action potential generation in a single compartment model. Lowering the temperature in this toy model slowed the rise of the action potential (Fig. 10A). The broadening of the action potential in these simulations is due to the temperature dependence of the voltage-gated potassium current we used. To quantify the change in action potential rise time, we numerically calculated the first derivative of the action potential (Fig. 10B) and plotted the maximal derivative as a function of temperature (Fig. 10C). This simulation predicted that the maximal rate of depolarization will be linearly dependent on the temperature (Fig. 10C). To test this prediction, we performed current-clamp experiment in neocortical pyramidal neurons over a range of temperatures (Fig. 10D). Since neocortical pyramidal neurons are very different than the single compartment model we did not expect that the experimental action potential waveform will resemble the simulated ones. However, the rise time of the experimental action potential was longer at lower temperatures (Fig. 10D) which was reflected in the first derivative (Fig. 10E). The maximal first derivative of the experimental action potential displayed a linear dependence on temperature (Fig. 10F) verifying the prediction of the single compartment simulation (Fig. 10C).

**Figure 9:**
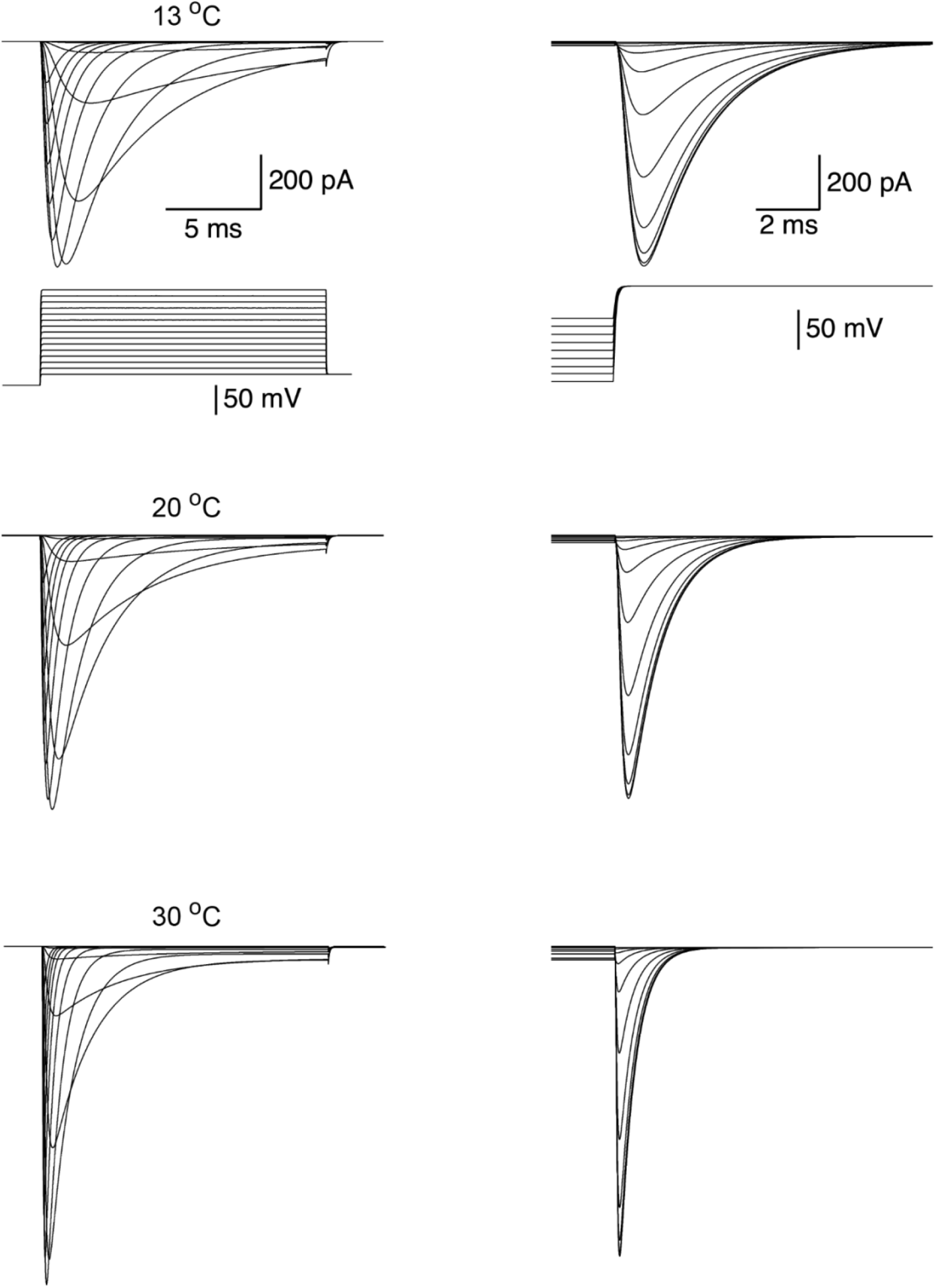
simulated voltage-clamp experiments of the sodium channel model as a function of temperature. Sodium currents simulated in response to an activation protocol (as in Figure 1) at four temperatures are shown on the left column. The voltage-clamp protocol is displayed below the top traces. Following a 100 ms pre-pulse to −110 mV the membrane potential was depolarized to the test potential at 10 mV increments. The right column displays currents simulated in response to an inactivation protocol as in Figure 5. The voltage-clamp protocol is displayed below the top traces. Following a 100 ms pre-pulse to several potentials at 10 mV increments the membrane potential was depolarized to the test potential of zero mV.

**Figure 10:**
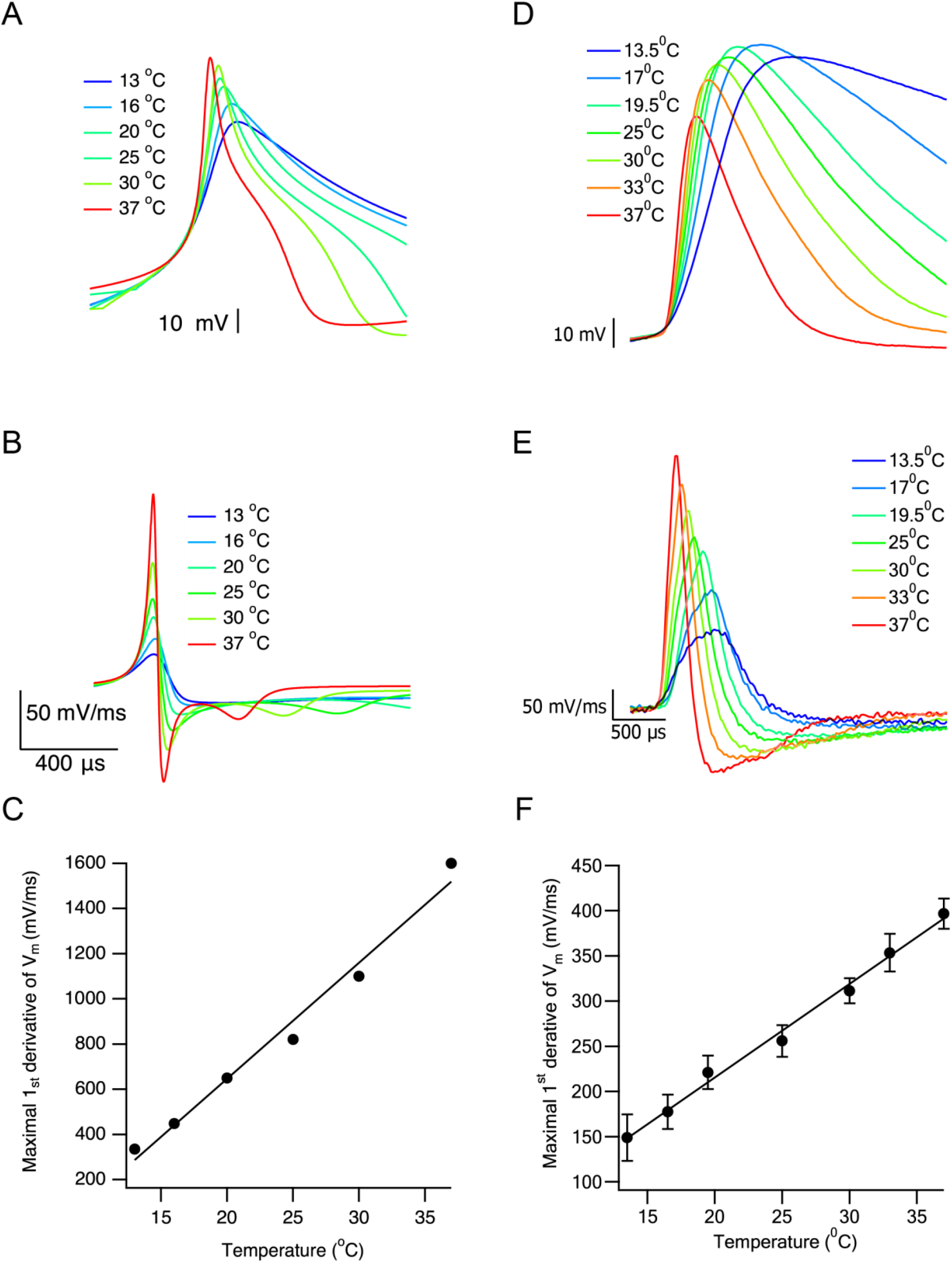
Simulated and experimental effect of temperature on action potential. A. Action potentials simulated in a single compartment model at various temperatures as indicated by the legend. B. numerical first derivative of the traces shown in A. C. The maximal first derivative calculated in B plotted as a function of temperature. Line is linear regression. D. Action potentials recorded in current clamp mode from Layer 5 neocortical pyramidal neurons at various temperatures as indicated by the legend. E. numerical first derivative of the traces shown in D. F. The maximal first derivative calculated in E plotted as a function of temperature. Line is linear regression. Error bars are SEM.

## Discussion

This study recorded sodium currents from nucleated patches extracted from the rat’s layer five neocortical pyramidal neurons at several temperatures from 13.5 to 30 °C. We use these recordings to model the kinetics of the voltage-gated sodium channel as a function of temperature using the Hodgkin-Huxley formalism and rate theory. We show that the temperature dependence of activation differs from that of inactivation. Furthermore, we show that the sustained current has a different temperature dependence than the fast current (Fig. 1). Our kinetic and thermodynamic analysis of the current provided a numerical model spanning the entire temperature range. This model reproduced vital features of channel activation and inactivation.

As expected, the fast inward sodium current and conductance increase with temperature. In addition, an increase in temperature results in a negative shift of the voltage dependence of activation, and a faster reaction kinetics, as was also demonstrated for Nav1.2 by other authors (Misra *et al.*, 2008; Hu *et al.*, 2009; Thomas *et al.*, 2009; Egri *et al.*, 2012). By contrast, the conductance of the sustained current decreases with temperature, and is estimated to be very small at physiological temperatures. Although the sustained sodium current is a small fraction of the transient current, it has a major effect on neuronal firing. A change of only a few percent in the sodium current may have a substantial effect on excitability (Stafstrom, 2007). Its significant role on repetitive firing was investigated extensively using experimental data and modeling (Kuo *et al.*, 2006; Stafstrom, 2007; Chadda *et al.*, 2017; Hsu *et al.*, 2018). The reduction of the sustained current as function of temperature is especially important since it indicates how modeling the channel at room temperature may lead to results that are irrelevant for biologically detailed models. This decrease in current as temperature increases suggests that, in the transition to the sustained configuration, the backward rate constant is more sensitive to temperature than the forward rate constant. Furthermore, from the temperature dependence of the channel (Figs. 2 and 6) it is clear that scanning the temperature range between 25°C and 30°C is not enough to provide a sufficient description of the temperature dependence of the rate constants. Some of the parameters appear not to change between 25°C and 30°C (for example in Figs. 2B and 2C). Cooling below 20°C is required to view subtle changes to the channel kinetics that are important once extrapolation to physiological temperatures is attempted. In the current study we recorded currents at four temperatures (Tab. 1). Our results suggest that more temperature points may be required to obtain a better estimation of the temperature dependence of the sodium channel.

Further thermodynamic analysis of the fast inward current indicates that the forward rate constant, *α*_*m*_, displays a higher sensitivity to temperature compared to the backward rate constant, *β*_*m*_ (Fig. 4). This is manifested in the shift of the activation curve with temperature (Fig. 1). Accordingly, the activation energy, Ea, is higher for the forward reaction. However, as for inactivation, that exhibits higher conductance with temperature increment, but with no change in voltage dependence, *α*_*h*_, and *α*_*h*_ display a similar temperature dependence (Fig. 8). Thus, the inactivation process is close to equilibrium at all tested membrane potentials and passively follows changes to the membrane potential. Correspondingly, there is a small difference between the activation energies of the forward and backward reactions (Fig. 8). In conclusion, the study demonstrated, under the assumptions of the Hodgkin-Huxley model that the primary energy barrier of channel gating resides in the activation process while the inactivation process is energetically close to equilibrium.

Many simulations express the thermal sensitivity of biological reactions using the temperature coefficient (Q_10_). This coefficient is defined as the ratio of two rate constants (a or b) separated by 10 degrees (Stover *et al.*, 1974). From our derivation of the rate constant for activation and inactivation we can derive Q_10_, for example for the forward rate of channel activation (Eq. 6).

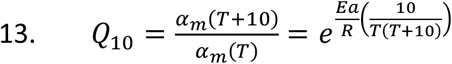

It is simple to show that Eq. 13 holds for all rate constants derived in this study, with the sole difference being that of the activation energy. However, plotting Q_10_ for all extracted rate constants (Fig. 11) demonstrates that this temperature coefficient is not constant for the voltage-gated sodium channel. Other authors also noted this property for sodium channels (Collins & Rojas, 1982; Thomas *et al.*, 2009) and mink channels (Allen & Mikala, 1998). Furthermore, Q_10_ was also shown to be voltage-dependent for neuronal sodium channels, Nav1.2, (Thomas *et al.*, 2009), and Kv channels (Ranjan *et al.*, 2019). Therefore, the practice of using a constant Q_10_ in numerical modeling of this channel may lead to inaccuracies. This warrants a revision of the way Q_10_ is used in numerical simulations. Furthermore, since *τ* = 1/(*α* + *β*) in the Hodgkin-Huxley model (with more complex expressions derived in Markov chain models), Q_10_ should be extracted directly from the rate constants and not from the time constant.

**Figure 11:**
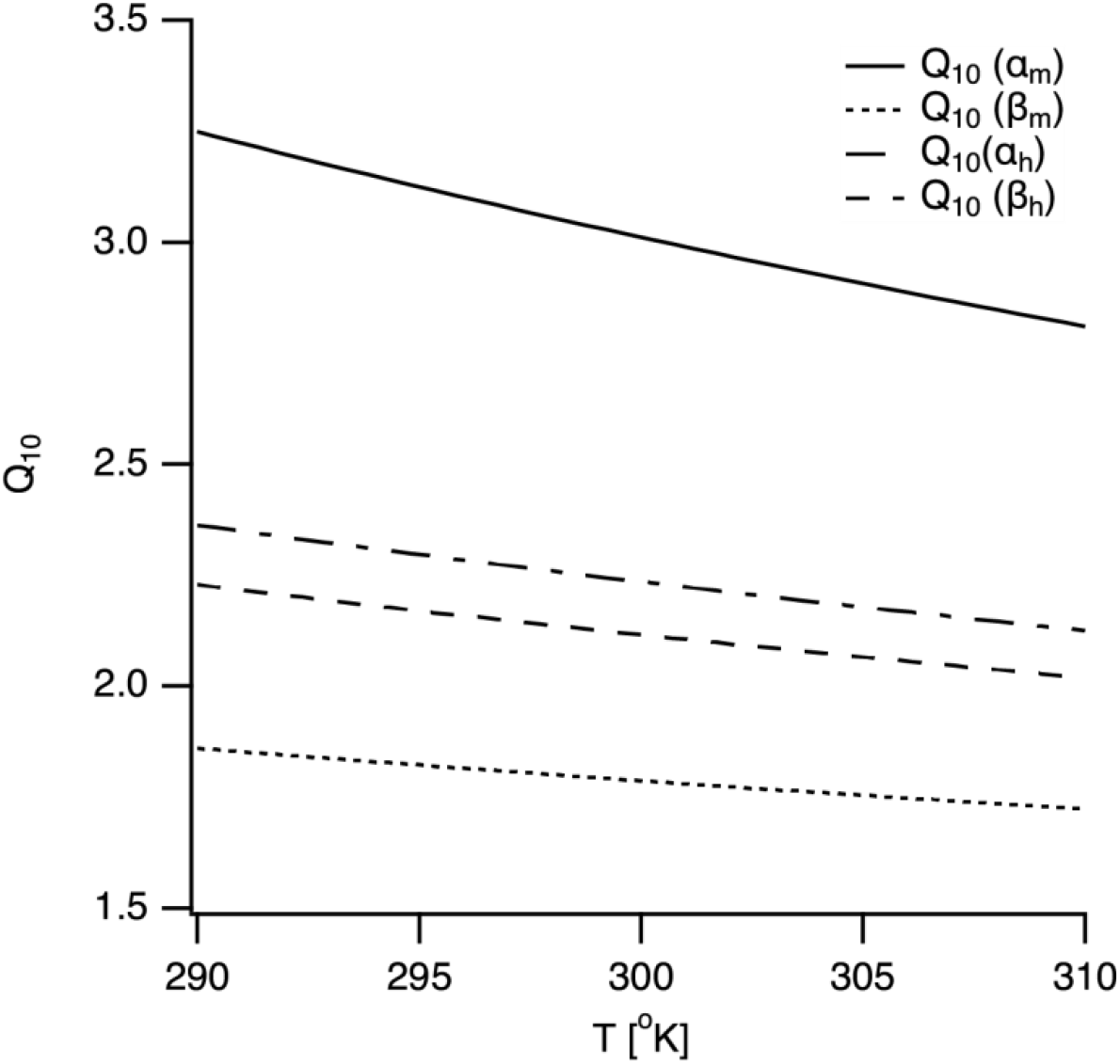
Temperature sensitivity of channel kinetics. The dependence of Q_10_ for each rate constant on temperature, calculated from eq. 13, is plotted as a function of temperature.

The numerical model we provide demonstrates the foundation and significance of models that include the changing effects of temperature on kinetic parameters. Our analysis demonstrates that it is possible to derive a single functional relationship describing the channel’s temperature and voltage dependence. However, the analysis presented in this paper has several limitations. To perform our analysis, we assumed that the channel obeyed a concerted gating model. Thus, the results of the analysis heavily depend on the selection of the model. The simplified model was able to reproduce the temperature dependence of the rising phase of the action potential. This simple sanity check demonstrated the validity of our description of the temperature dependence. However, a more detailed model for channel kinetics is required to capture the complex firing of cortical pyramidal neurons fully. Such a model would also have to take into account the kinetics of the sustained current, which we decided to neglect in the analysis. For the sake of our limited analysis, this was a reasonable assumption, especially since the data suggested that the sustained current should be small or even negligible at physiological temperatures (Fig. 2A). The increase in peak sodium conductance and the decrease in sustained conductance as a function of temperature may suggest a temperature-dependent change in the channel configuration, gradually shifting it from fast to sustained activity. Considering that in the cortex Nav1.2 channels rarely experience low temperatures, the function of this possible mode shifting is not clear. Such a mechanism may be of greater importance in peripheral nerves exposed to more significant temperature shifts.

## Acknowledgments

This work was supported by grants from the Israel Science Foundation to AK (#225/20).

## Author contribution

AK and MA designed the study, performed the experiments, and analyzed the data. NDK and AK drafted and revised the manuscript. All authors read and approved the final version of the manuscript for publication.

## Data accessibility

Data will be available upon a request to corresponding author. Model code is available from ModelDB (note to reviewers: model code will be deposited on acceptance of article).

